# LAND: Latent Aligned Neural-Behavioral Dynamics via Flow Matching for Generalizable Movement Decoding

**DOI:** 10.64898/2026.08.19.745390

**Authors:** Ruwei Yao, Junze Zheng, Yanlu Wang, Wenzheng Li, Xiaolong Zou, Bo Hong

**Affiliations:** Tsinghua University, Beijing, China

## Abstract

Generalizable movement decoding remains a central challenge for invasive brain–computer interfaces (BCIs), as decoders trained under limited calibration conditions often fail to generalize to unseen movement speeds, limbs, and subjects. Existing decoding methods are typically trained on paired data collected under restricted conditions. How to incorporate behavioral structure from unpaired data for robust out-of-distribution (OOD) decoding therefore remains unresolved. To address this, we propose **LAND** (**L**atent **A**ligned **N**eural-behavioral **D**ynamics), a framework that aligns latent neural and behavioral dynamics through flow matching. By learning a neural-to-behavioral transport map and using behavioral-dynamics priors from unpaired data to encourage structured neural manifolds, LAND regularizes representation geometry to promote cross-domain generalization. We evaluate LAND on synthetic neural data, epidural BCI recordings from a tetraplegia participant, and multi-electrode array (MEA) recordings from nonhuman primates (NHPs). Across these settings, LAND improves zero-shot generalization to OOD movement speeds and yields speed-modulated manifolds. With limited target-domain fine-tuning, it further improves transfer across limbs and subjects. These results support flow-based neural–behavioral alignment with unpaired kinematic priors as an approach for learning transferable neural representations and robust movement decoding across behavioral and recording domains.

## Introduction

Generalizable decoding of motor behavior from neural activity remains challenging for the clinical translation of BCIs (Li et al. 2026). These systems must map high-dimensional, noisy cortical signals to diverse behavioral trajectories (Lorach et al. 2023), thereby restoring movement for neurologically impaired individuals (Liu et al. 2024). However, real-world BCI deployment often entails distribution shifts, such as movement speeds and rhythms that differ from those observed during training (Yao et al. 2025), as well as transfer to a new limb (Merrick et al. 2022) or subject (Li et al. 2026). This mismatch between limited training distributions

Continuous BCI decoders model neural–behavioral relationships using kinematic priors, deep temporal networks, or Neural Dynamics Models (NDMs) (Sani et al. 2021; Wu et al. 2002; Kao et al. 2014; Sussillo et al. 2016; Sun et al. 2024; Glaser et al. 2020). NDMs exploit the hypothesis that neural activity is governed by low-dimensional dynamical systems (Vyas et al. 2020; Gallego et al. 2017; Churchland et al. 2012) and can capture intrinsic neural manifolds (Pandarinath et al. 2018; Karpowicz et al. 2025). Nevertheless, high-capacity decoders can overfit restricted training conditions, while behavior-guided dynamical models generally rely on paired data from the same domain. Bridging accurate neural-behavior modeling with transferable representation learning therefore remains an open challenge for BCI (Xu et al. 2021).

Behavioral kinematics offer an accessible proxy for the latent dynamics underlying neural motor control. Unlike neural recordings, behavioral data can be acquired at much larger scales and naturally encompass diverse movement conditions (Yao et al. 2025). Since motor behavior emerges from structured low-dimensional neural dynamics (Mathis and Mathis 2025; Schneider, Lee, and Mathis 2023), behavioral dynamics are expected to preserve transferable task structure across domains. This motivates leveraging behavioral dynamics priors to align neural representations, promoting domain-invariant latent dynamics instead of fitting domain-specific patterns. However, enforcing point-to-point embedding level alignment risks noise overfitting and geometric distortion. Flow matching (Lipman et al. 2023; Liu, Gong, and Liu 2022) can overcome this by learning a continuous velocity field for flexible, distribution-level alignment.

Here we propose **LAND** (**L**atent **A**ligned **N**euralbehavioral **D**ynamics), a generalizable movement decoding framework. LAND leverages accessible unpaired behavioral data to learn broader kinematic priors and shape a structured manifold. The framework learns neural-to-behavioral transport via flow matching and aligns neural representations with that manifold. By emphasizing dynamical structure shared across domains, this formulation aims to produce representations that transfer across speeds, homologous limbs, and subjects (Figure 1) while supporting inference using neural activity alone. We evaluate LAND on both synthetic data and human/NHPs BCI datasets spanning epidural field potentials and single-unit recordings.

**Figure 1:**
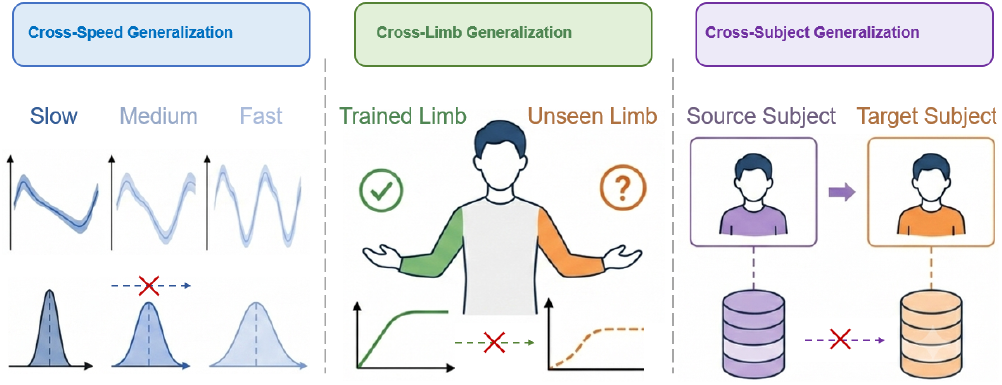
Cross-domain generalization challenges in movement decoding. From left to right: generalization across unseen speeds, limbs, and subjects. Red crosses denote domain shifts beyond direct source-domain training. and unseen OOD conditions impairs robust and scalable decoding.

Our main contributions are summarized as follows:

- **Latent Space Flow Matching:** We propose LAND, which learns neural-to-behavioral transport through flow matching and aligns latent neural and behavioral dynamics for movement decoding.
- **Structured Neural Representations:** LAND uses behavioral kinematic priors learned from unpaired data to encourage structured neural manifolds that generalize across speeds, limbs, and subjects.
- **Cross-Domain Evaluation:** Across synthetic, epidural BCI, and NHP single-unit recordings, LAND improves zero-shot cross-speed decoding and, with limited finetuning, transfer across limbs and subjects.

## Related Work

### Neural Dynamics Models

Classical neural decoding predominantly relies on end-to-end “black-box” sequence models—such as RNNs (Xie, Schwartz, and Prasad 2018), TCNs (Rabbani et al. 2025), Transformers (Komeiji et al. 2024), and State Space Models (Suzuki et al. 2025). Although effective at modeling complex temporal dependencies, these “black-box” mappings (Glaser et al. 2020) do not explicitly identify or constrain neural dynamics, creating severe bottlenecks in generalizing to OOD scenarios. Grounded in neural population dynamics (Gallego et al. 2017; Shenoy, Sahani, and Churchland 2013), Neural Dynamics Models (NDMs) infer low-dimensional latent manifolds to enable robust estimation and generalizable representations (Perkins et al. 2025). Within NDMs, generative frameworks (e.g. LFADS (Pandarinath et al. 2018)) extract latent factors via neural auto-encoding (Keshtkaran et al. 2022; Sedler, Versteeg, and Pandarinath 2023) or diffusion models (Kapoor et al. 2024), but may risk discarding behaviorally relevant features (Mathis and Mathis 2025). Conversely, behavior-guided NDMs like PSID (Sani et al. 2021), DPAD (Sani, Pesaran, and Shanechi 2024) incorporate behavioral supervision to capture behavior-related dynamics. Yet these approaches rely on paired observations in standard formulations; the behavioral supervision is therefore limited.

### Neural Foundation Models

Recent advances in Neural Foundation Models have expanded data scale and cross-context generalization through large-scale pretraining. Models such as STNDT (Le and Shlizerman 2022) or NDT2 (Ye et al. 2023) focus on spatiotemporal neural sequence modeling and multi-context pretraining. POYO (Azabou et al. 2023; Ryoo et al. 2026) introduces cross-dataset, multi-task supervised neural decoding, while frameworks like NEDS (Zhang et al. 2025) employ multimodal masked cross-reconstruction for joint representation learning. These foundation models broaden task coverage and facilitate cross-session and cross-subject transfer by learning reusable neural representations or cross-modal predictive features. Complementarily, LAND introduces an explicit behavioral dynamics prior from unconstrained, unpaired behavioral data to promote the transfer of behavioral geometry across domain shifts.

### Neural–Behavior Alignment

A parallel line of research learns joint neural–behavioral representations for robust and interpretable decoding (Mathis and Mathis 2025). Contrastive methods construct positive and negative pairs to learn discriminative representations; for example, CEBRA (Schneider, Lee, and Mathis 2023) associates neural activity with behavioral variables or time, and contrastive objectives have also been used for visual–neural multimodal alignment (Song et al. 2023). PNBA (Zhu et al. 2025) employs probabilistic matching objectives to construct aligned representations, focusing on addressing session and subject variability. Separately, flow matching and rectified flows learn continuous transport vector fields between distributions (Lipman et al. 2023; Liu, Gong, and Liu 2022). LAND combines these perspectives by learning a neural-to-behavior latent transport from paired observations while using auxiliary unpaired kinematic trajectories to shape the behavioral latent space. Rather than enforcing only samplewise correspondence, LAND transfers neural embeddings to a behavioral representation with broader temporal variation, while retaining neural-only decoding at inference.

## Methods

To address the aforementioned challenges in neural decoding generalization and the limitations of existing approaches, we propose the LAND framework.

### Problem Formulation

Let **y** ∈ *Y* ⊂ ℝ^*T* ×*C*^ and **x** ∈ *X* ⊂ ℝ^*T* ×*N*^ denote neural activity and behavioral kinematic sequences, respectively, where *T* is the sequence length and *C* and *N* are the neural-feature and behavioral dimensions. The fundamental BCI problem is to learn a causal mapping *D* : *Y* → *X* that remains effective under shifts in movement speed, limb, and subject. In practice, calibration pairs (**y**_pair_, **x**_pair_) are acquired in restricted source domains. The resulting decoder must therefore generalize without target neural data to unseen movement speeds and adapt with limited target-domain calibration to new limbs or subjects.

### Framework Architectures

LAND comprises a neural encoder *f*_n_, a behavioral encoder *f*_b_, a flow-alignment network *v*_*θ*_, and a shared behavioral decoder *D*_*ψ*_ (Figure 2a). For a paired sequence, the encoders produce time-resolved embeddings

**Figure 2:**
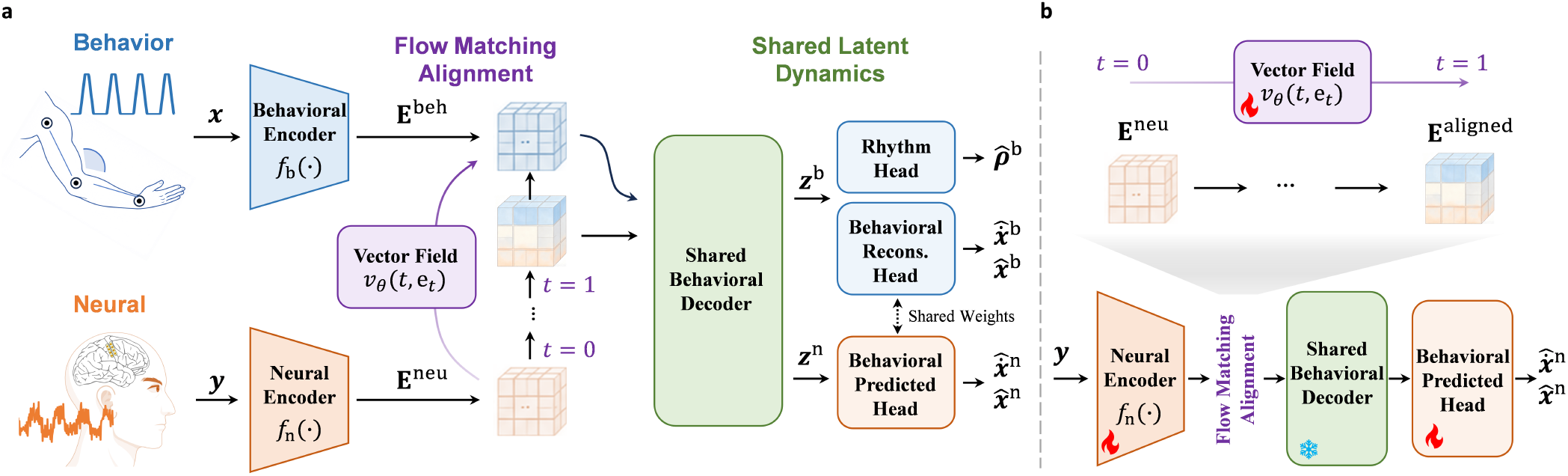
Overview and cross-domain adaptation of LAND. **a.** Behavioral kinematics **x** and neural observations **y** are encoded as **E**^beh^ and **E**^neu^. The vector field *v*_*θ*_(*t*, **e**_*t*_) transports neural embeddings to **E**^aligned^, and the shared behavioral decoder produces behavioral and neural factors. The behavioral reconstruction and neural prediction heads each output kinematics and velocities; the behavioral path additionally reconstructs rhythm. **b**. For cross-limb and cross-subject transfer, we fine-tune the neural encoder, flow field, and behavioral prediction head (fire symbols), while freezing the shared behavioral decoder (snowflake).

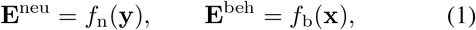

where **E**^neu^, **E**^beh^ ∈ ℝ^*T* ×*M*^ and *M* is the embedding dimension. The neural encoder is a multilayer perceptron (MLP) applied independently at each time point. The behavioral encoder is a unidirectional GRU followed by a linear projection, which encodes the position components of the kinematic input. It is trained with both paired kinematics and auxiliary unpaired trajectories, acquired from natural behavior through pose estimation or generated by randomized simulations, to learn a manifold spanning a broader behavioral prior. These heterogeneous encoders allow neural activity and behavior to retain modality-specific representations before alignment. Flow alignment maps the neural embedding onto this behavioral embedding without requiring the heterogeneous embeddings to coincide pointwise. It defines a time-dependent vector field *v*_*θ*_(*t*, **e**_*t*_) over the embedding space. The field is parameterized by a multilayer perceptron that receives an embedding and a sinusoidal representation of *t* ∈ [0, 1]. Starting from a neural embedding, we obtain its behavior-aligned counterpart by integrating the learned ordinary differential equation

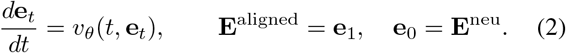

We approximate this transport with a fixed-step forward Euler solver (10 steps in our implementation). Thus, the neural pathway is **y E**^neu^ **E**^aligned^, whereas the behavioral pathway directly uses **E**^beh^.

A shared causal decoder extracts latent dynamic factors from both paths,

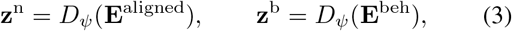

followed by a behavioral reconstruction head and rhythm head. The behavioral reconstruction head jointly outputs re-constructed kinematics 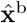 and their velocities 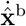, while the rhythm head outputs 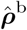. Likewise, the behavioral prediction head jointly outputs predicted kinematics 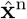 and velocities 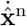. The decoder can be instantiated with different temporal-model backbones, such as GRUs and TCNs; all variants operate causally. Sharing this decoder makes behavioral temporal structure available to the transported neural representations without assuming that raw neural and behavioral embeddings should coincide.

The framework has two complementary training paths (Figure 2a). The behavioral path processes both paired kinematics and unpaired auxiliary trajectories, learning to reconstruct kinematics and their velocities while predicting rhythm. The neural path encodes paired neural observations, transports their embeddings through the flow field, and decodes the resulting factors to kinematics and velocities. During online inference, only this neural path is retained: the model integrates the vector field from each neural embedding and predicts the trajectory and velocity using the shared behavioral decoder and behavioral prediction head, without behavioral or rhythm inputs.

#### Cross-domain fine-tuning

For a new homologous limb or subject, we initialize LAND from a source-domain model and use limited target-domain paired data to adapt the neural interface (Figure 2b). Because the shared behavioral decoder encodes task dynamics that should be invariant across domains, whereas neural feature distributions and their alignment can change substantially, we freeze the decoder and optimize the target neural encoder *f*_n_, flow field *v*_*θ*_, and behavioral prediction head with *L*_dec_. This parameter-efficient adaptation preserves the learned behavioral manifold and its temporal dynamics while calibrating the target-specific neural interface.

### Objective

LAND jointly uses paired data to learn the neural-to-behavioral correspondence and unpaired kinematics to expand the behavioral prior. For a corresponding pair of embedding vectors **e**_0_ ∈ **E**^neu^ and **e**_1_ ∈ **E**^beh^ at the same time index, we sample *t* ∼ *U* (0, 1) and form the linear probability path

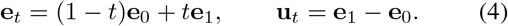

The flow-matching objective regresses the vector field to the constant velocity of this path:

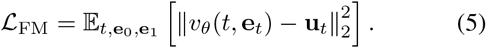

Gradients from this objective update the neural encoder and flow network, while the behavioral target embedding is stopgradient. This asymmetric construction anchors the learned transport to the behavioral manifold rather than permitting both manifolds to drift together.

#### Paired-data objectives

After flow integration, we encourage the decoded neural factors to match the behavioral factors from the same paired trial,

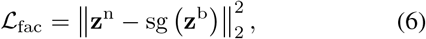

where sg(·) denotes stop-gradient. Direct supervision of kinematics and their velocities ensures that the transported representation remains useful for decoding,

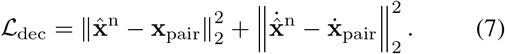

Unlike direct embedding matching, *L*_FM_ learns a trajectory-dependent transformation between the two distributions, and *L*_fac_ applies consistency after this transport.

#### Behavioral-prior objectives

Let 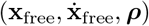 be an unpaired kinematic sequence, its velocity target, and its rhythm target. Training on these trajectories extends the behavioral manifold beyond the restricted conditions represented by paired neural data. The behavioral pathway reconstructs kinematics and velocities while predicting rhythm,

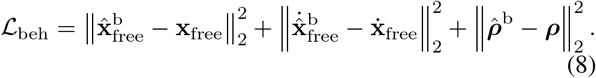

To prevent a low-rank or redundant behavioral factor space, we additionally regularize the factors **Z**^b^ ∈ ℝ^*K*×*L*^ across all batch-time samples. With covariance **C** of mean-centered factors, the diversity loss penalizes insufficient marginal variance and cross-factor covariance:

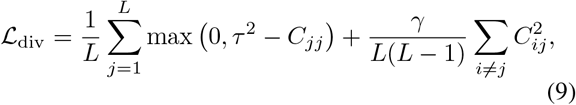

where *τ* is a variance threshold and *γ* controls the covariance penalty.

The final training objective is

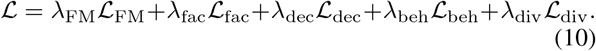

## Experiments and Results

We evaluated LAND on synthetic neural decoding data, epidural BCI recordings from a tetraplegia participant, and MEA recordings from two NHPs. All models in this study were trained on a workstation equipped with an Intel Xeon Gold 5218R CPU and an NVIDIA GeForce RTX 3090 GPU, using Python 3.10 and PyTorch. LAND improved generalization to OOD movement speeds and learned interpretable, speed-modulated neural–behavioral representations. We further evaluated limited-calibration transfer across limbs and subjects while preserving the learned behavioral dynamics for causal neural decoding.

### Synthetic Dataset Validation

#### Synthetic Dataset and Comparison

We constructed a synthetic neural decoding dataset to intuitively examine its properties. Limb trajectories (*N* = 3) were composed of trapezoidal gestures. A naive nonlinear recurrent dynamics model generated the corresponding neural activity (*C* = 8) from its recurrent state and fixed 8 × 3 position and velocity interaction mappings; dominant dimension-specific weights and weaker cross-dimensional weights produced mixed neural tuning, with additive process and observation noise. Paired training data followed one fixed rhythm, whereas the test set contained three individual movements at five rhythms spanning slower and faster conditions on both sides of the training distribution. We compared LAND, which additionally learned from unpaired free-timing trajectories through its behavioral branch, with an architecturally matched End-to-End decoder trained only on paired data (details in the Supplementary).

#### OOD Generalization

The End-to-End baseline initially improved but then overfit the fixed training rhythm, causing test error to rebound as training continued. In contrast, LAND continued to improve and outperformed the optimally early-stopped baseline (MSE = 0.025 ± 0.009 vs. 0.033 ± 0.017; Welch’s *t*-test, *p* = 5.2 × 10^−3^; Figure 3a). Narrowing the auxiliary temporal distribution from width 2 to 0 progressively degraded generalization (Figure 3b); at width 0, trajectory MSE increased by 41% (*p* = 4 × 10^−4^) and rhythm error by 71% (*p* = 0.023). This result attributes LAND’s extrapolation to the temporal diversity of unpaired kinematics rather than merely to the presence of an additional branch. Qualitative trajectories are provided in the Supplementary.

**Figure 3:**
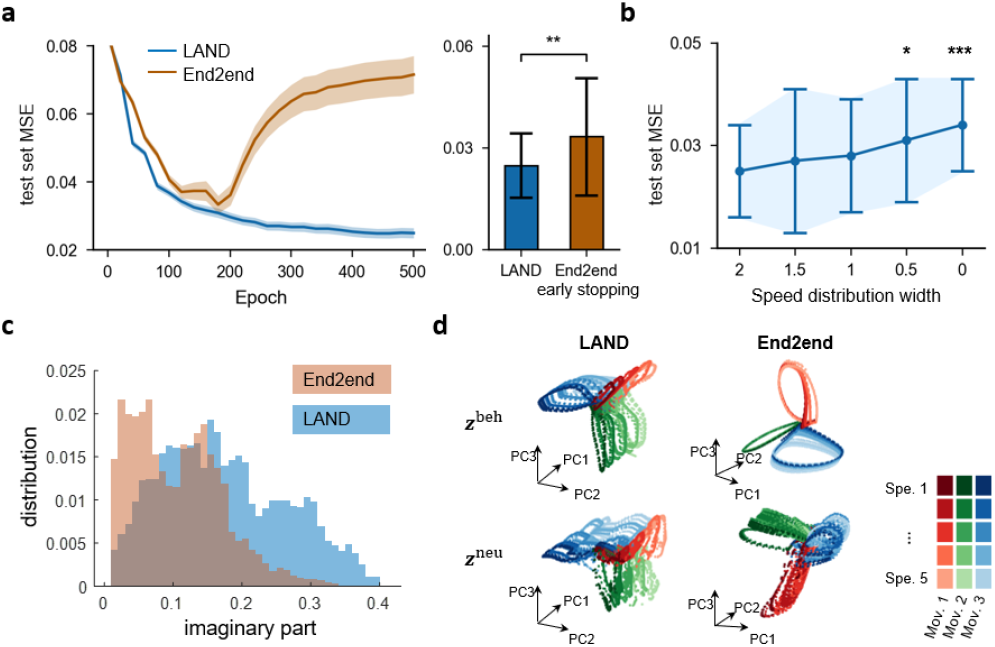
Synthetic-data validation of LAND. **a.** Test MSE across training epochs (left) and LAND versus the optimally early-stopped End-to-End baseline (right). Shading denotes SE; error bars denote SD. **b**. LAND test MSE as the temporal-distribution width of auxiliary data decreases from 2 to 0. Error bars denote SD, and asterisks mark significant differences from width 2. **c**. Distributions of the imaginary components of decoder-Jacobian eigenvalues. **d**. PCA projections of behavioral factors **z**^beh^ (top) and neural factors **z**^neu^ (bottom) for LAND and End-to-End. Colors denote the 3 movement dimensions and shade intensity denotes the 5 speed levels.

#### Learned Dynamics and Manifolds

To elucidate the dynamical mechanisms underlying LAND, we analyzed the Jacobian matrices of the decoder (details in the Supplementary). LAND learned a broader distribution of decoder-Jacobian imaginary eigenvalues than End-to-End (Levene’s test, *p <* 10^−5^; Figure 3c), consistent with a wider range of rotational frequencies available to the recurrent decoder. Its behavioral and aligned neural factors also separated movement dimensions while varying systematically with speed, including OOD rhythms; notably, this organization was retained in neural factors despite paired neural data being observed at only one rhythm. End-to-End factors were less coherently organized (Figure 3d). Thus, behavioral alignment transferred reusable speed-modulated geometry to the neural representation.

### Epidural BCI Validation

#### Clinical study and paradigm

We evaluated LAND with a wireless minimally invasive BCI. Its coin-sized processor is embedded in the skull, and two four-contact arrays (8 channels in total) record epidural EEG over the right primary motor and somatosensory cortices. The participant was a man with tetraplegia caused by a complete C4 spinal cord injury (AIS-A), enrolled in an ethics-approved clinical trial of an implantable closed-loop BCI. He performed video-guided flexion and extension of the left (contralateral) and right (ipsilateral) elbows at *Fast, Middle*, and *Slow* speeds (Figure 4a). Each trial comprised rest, flexion, hold, and extension phases, with speed controlled by the movement-phase duration (details in the Supplementary).

**Figure 4:**
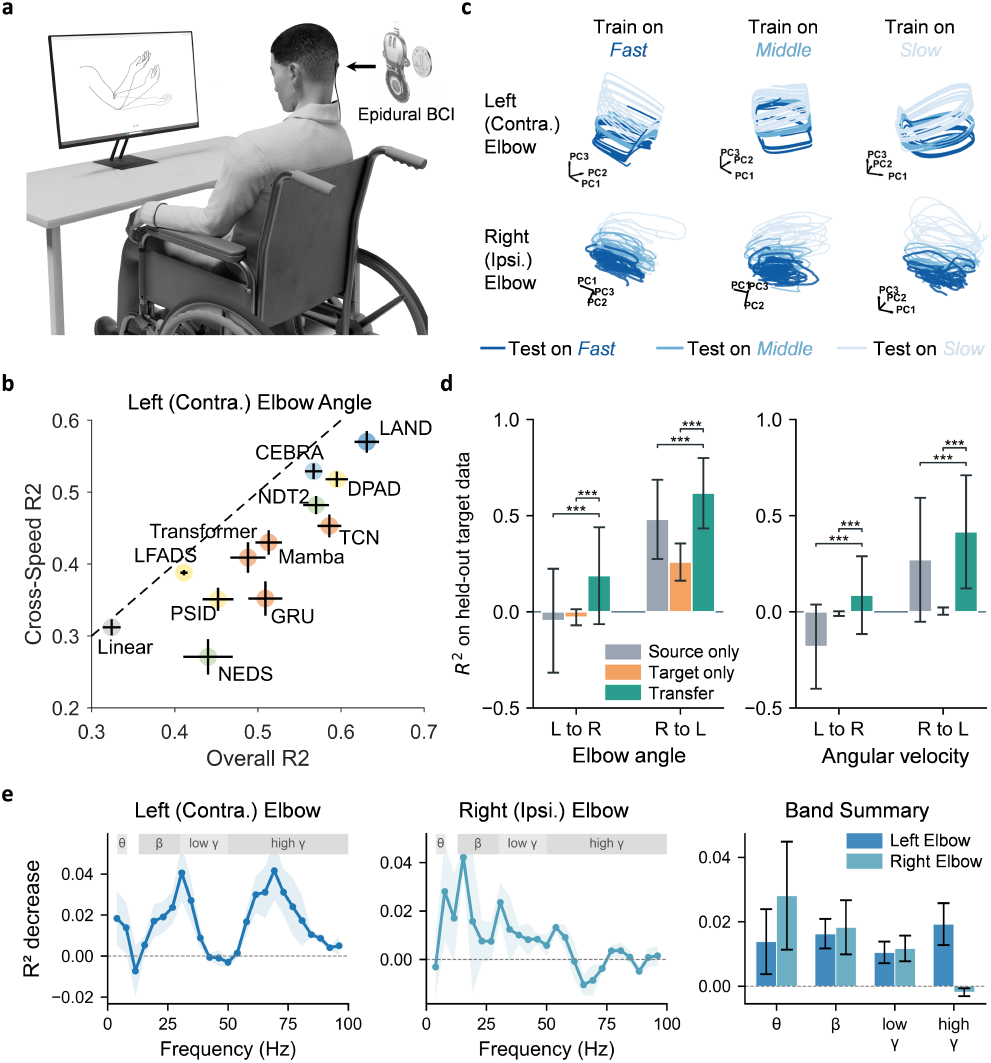
Epidural BCI validation. **a.** Video-guided bilateral elbow tracking with the epidural BCI. **b**. Overall versus cross-speed *R*^2^ for left-elbow angle decoding; points above the diagonal generalize better across speeds than they decode on average. **c**. PCA projections of LAND latent neural dynamics for the contralateral left and ipsilateral right elbows. Columns denote the training speed, and line colors denote the testing speed. **d**. Cross-limb transfer from left to right (L to R) and right to left (R to L) for elbow angle and angular velocity. Bars show held-out target-domain *R*^2^, error bars indicate variability, and brackets mark significant pairwise differences. **e**. Frequency-wise perturbation sensitivity, measured as the *R*^2^ decrease and averaged across angle and velocity decoding.

#### Processing, models, and evaluation

Preprocessed epidural EEG signals were converted into PSD feature sequences. Both elbow angle and angular velocity (*ω*) were decoded. We used a GRU behavioral decoder for LAND, trained with randomized-speed auxiliary trajectories, and a causal shared decoder. Comparisons included linear decoders, direct GRU, TCN, Transformer, Mamba, NDMs (LFADS, PSID, DPAD), contrastive models (CEBRA), and neural foundation models (NDT2, POYO, NEDS). For each limb and output, models were trained separately on each speed in a 3 × 3 train–test design. We report overall *R*^2^ across all nine train–test combinations and cross-speed *R*^2^ across the six off-diagonal combinations; separate intra-speed summary statistics are not reported. Implementation and evaluation details are provided in the Supplementary.

#### Bilateral decoding and cross-speed generalization

The results reveal a marked hemispheric asymmetry (Table 1). Under speed shifts, LAND attained the strongest contralateral left-elbow decoding, reaching *R*^2^ = 0.57 for angle and 0.38 for velocity. These values exceed the strongest non-LAND baselines (CEBRA for angle and DPAD for velocity) by 8% and 19%, respectively. The Overall–Cross comparison in Figure 4b further indicates that this advantage persists relative to average decoding accuracy. In contrast, ipsilateral right-elbow decoding was substantially harder for every method: LAND reached cross-speed *R*^2^ values of 0.10 for angle and 0.04 for velocity. Thus, the clearest gains occur for contralateral decoding, and ipsilateral velocity remains an important limitation (see Supplementary for detailed data).

**Table 1:** Epidural BCI cross-speed decoding *R*^2^, mean ± SD. TF: Transformer. Results average the off-diagonal train–test combinations.

| Method | Left elbow (Contralateral) |  | Right elbow (Ipsilateral) |  |
| --- | --- | --- | --- | --- |
|  | Angle | Velocity | Angle | Velocity |
| Linear | .31 ± .12 | .00 ± .08 | -.01 ± .09 | -.03 ± .04 |
| GRU | .35 ± .26 | -.03 ± .34 | -.06 ± .39 | -.18 ± .34 |
| TCN | .45 ± .18 | .21 ± .21 | .09 ± .32 | -.07 ± .26 |
| TF | .41 ± .23 | .13 ± .28 | -.02 ± .04 | -.01 ± .02 |
| Mamba | .43 ± .19 | .25 ± .22 | .02 ± .14 | -.04 ± .09 |
| LFADS | .39 ± .04 | .08 ± .10 | .00 ± .05 | -.01 ± .01 |
| PSID | .35 ± .18 | .09 ± .35 | .08 ± .07 | -.09 ± .20 |
| DPAD | .52 ± .12 | .32 ± .22 | .09 ± .06 | .00 ± .02 |
| CEBRA | .53 ± .12 | .00 ± .03 | -.05 ± .14 | -.02 ± .02 |
| NDT2 | .48 ± .14 | .26 ± .27 | -.01 ± .04 | -.08 ± .12 |
| POYO | -.01 ± .28 | -.22 ± .47 | -.13 ± .17 | -.29 ± .42 |
| NEDS | .27 ± .27 | -.04 ± .53 | -.01 ± .03 | -.02 ± .03 |
| **LAND** | **.57 ± .17** | **.38 ± .22** | **.10 ± .33** | **.04 ± .15** |

#### Structured speed manifolds

Across models trained on fast, middle, or slow movements, PCA projections of LAND factors formed organized rotational trajectories for both elbows (Figure 4c). The trajectories for the three testing speeds occupied related paths within each training-speed condition, including unseen-speed tests, whereas the ipsilateral geometry was visibly less compact. Together with the cross-speed results, this organization suggests that alignment transfers movement-phase and speed structure into the neural representation rather than relying only on speed-specific temporal templates (Saxena et al. 2022; Churchland and Shenoy 2024). **Cross-limb generalization**. We initialized the decoder from one elbow and adapted it to the other while preserving shared behavioral dynamics (Figure 2b). For fair comparison, both target-only and transfer used 10% of target-domain paired data; source pretraining and target adaptation included all three speeds. Table 2 reports held-out transfer *R*^2^. In the challenging L → R direction, LAND was best for both angle (0.19±0.25) and velocity (0.09±0.20), improving on the next strongest baseline, PSID, by 0.08 and 0.06 *R*^2^, respectively. In the reverse R→L direction, LAND achieved 0.62 ± 0.18 for angle and 0.42 ± 0.29 for velocity, second to NDT2 (0.67 and 0.49). This asymmetry reflects the ipsilateral right-elbow target in L→R transfer, which is harder to decode than the contralateral left-elbow target in R L transfer. Thus, LAND provides a useful initialization for limited-calibration adaptation across limbs (see Supplementary).

**Table 2:** Epidural BCI cross-limb transfer *R*^2^, mean ± SD.

| Method | L→R |  | R→L |  |
| --- | --- | --- | --- | --- |
|  | Angle | Velocity | Angle | Velocity |
| LFADS | 0.04 ± 0.21 | 0.01 ± 0.12 | 0.33 ± 0.14 | 0.05 ± 0.10 |
| PSID | 0.11 ± 0.17 | 0.03 ± 0.15 | 0.43 ± 0.14 | 0.28 ± 0.18 |
| DPAD | -0.19 ± 0.53 | -0.48 ± 0.60 | 0.52 ± 0.24 | 0.25 ± 0.34 |
| NDT2 | 0.10 ± 0.10 | 0.02 ± 0.08 | **0.67 ± 0.07** | **0.49 ± 0.13** |
| POYO | 0.03 ± 0.05 | 0.01 ± 0.06 | 0.02 ± 0.02 | 0.03 ± 0.02 |
| NEDS | 0.08 ± 0.10 | 0.01 ± 0.09 | 0.56 ± 0.12 | 0.34 ± 0.18 |
| **LAND** | **0.19 ± 0.25** | **0.09 ± 0.20** | 0.62 ± 0.18 | 0.42 ± 0.29 |

#### Physiological plausibility

Frequency-wise perturbation analysis, averaged across angle and velocity decoding, revealed dissociable spectral contributions (Figure 4e). Contralateral decoding was most sensitive to high-*γ* activity (55– 80 Hz) and *β*-band (15–30 Hz), whereas the lower-SNR ipsilateral signals showed greater reliance on *β*-band ERD. This perturbation profile is consistent with established movement-related spectral patterns (Miller et al. 2007; Manning et al. 2009; Pfurtscheller and Da Silva 1999; Jurkiewicz et al. 2006; Shibasaki et al. 1980). See Supplementary for details.

### NHP Dataset Validation

We further evaluated LAND on the upper-limb cycling dataset of Saxena et al. (2022), which contains single-unit recordings and continuous limb kinematics from two NHPs, monkey C (74 neural channels) and monkey D (52 channels; Figure 5a). For each monkey, we used nine forward-cycling speed conditions and decoded normalized horizontal position and velocity. We evaluated three cross-speed splits by training on one adjacent speed triplet (levels 1–3, 4–6, or 7– 9) and testing on the remaining six levels, thereby assessing generalization to unseen movement rates rather than random temporal holdout. We configured a GRU as the behavioral decoder for LAND, and simulated auxiliary behavioral data with diverse rhythms (details in the Supplementary).

**Figure 5:**
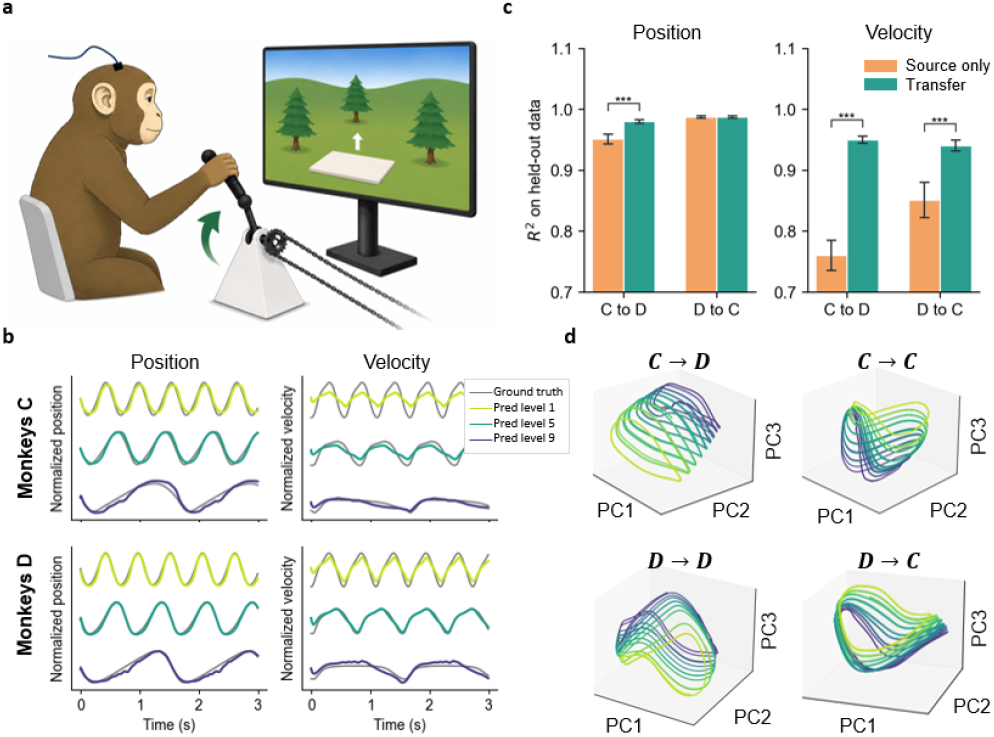
NHP cycling validation. **a.** Monkeys performed upper-limb cycling at nine target speeds (adapted from (Saxena et al. 2022)). **b**. LAND predictions and ground truth. Model trained on speed triplet 4–6. **c**. Held-out target-subject decoding after transfer versus target-only training. **d**. PCA projections of decoder latent dynamics before and after cross-subject fine-tuning.

#### Cross-speed decoding

LAND achieved the strongest position decoding for both monkey C (*R*^2^ = 0.946 ± 0.040) and monkey D (*R*^2^ = 0.978 ± 0.003; Table 3). Relative to the strongest non-LAND position baselines, these correspond to gains of approximately 2% for monkey C and 3% for monkey D. LAND also achieved the strongest velocity decoding for monkey C (0.730 ± 0.266), whereas POYO achieved the highest velocity *R*^2^ for monkey D. Across low, intermediate, and high held-out speeds, LAND predictions closely followed the ground-truth position and velocity traces (Figure 5b), preserving cycle phase, trajectory amplitude, and frequency beyond the speed conditions used for neural calibration. This behavior indicates that the learned representation captures cycling dynamics rather than merely interpolating among the calibrated rates (see Supplementary for detailed data).

**Table 3:** NHPs cross-speed decoding *R*^2^, mean ± SD. TF: Transformer. All entries evaluate held-out speed conditions.

| Method | Monkey C |  | Monkey D |  |
| --- | --- | --- | --- | --- |
|  | Position | Velocity | Position | Velocity |
| Linear | 0.89 ± 0.08 | 0.68 ± 0.30 | 0.90 ± 0.06 | 0.78 ± 0.15 |
| GRU | 0.92 ± 0.07 | 0.70 ± 0.29 | 0.95 ± 0.04 | 0.65 ± 0.33 |
| TCN | 0.93 ± 0.05 | 0.71 ± 0.26 | 0.93 ± 0.05 | 0.73 ± 0.23 |
| TF | 0.86 ± 0.12 | 0.31 ± 0.68 | 0.90 ± 0.06 | 0.26 ± 0.71 |
| Mamba | 0.64 ± 0.09 | 0.15 ± 0.22 | 0.56 ± 0.07 | 0.47 ± 0.14 |
| LFADS | 0.36 ± 0.34 | 0.16 ± 0.46 | 0.89 ± 0.07 | 0.65 ± 0.24 |
| PSID | 0.58 ± 0.32 | 0.37 ± 0.45 | 0.88 ± 0.01 | 0.60 ± 0.34 |
| DPAD | 0.81 ± 0.03 | 0.64 ± 0.07 | 0.92 ± 0.04 | 0.75 ± 0.11 |
| CEBRA | 0.75 ± 0.27 | -0.29 ± 2.34 | 0.81 ± 0.18 | 0.27 ± 1.09 |
| NDT2 | 0.90 ± 0.03 | 0.71 ± 0.11 | 0.94 ± 0.04 | 0.84 ± 0.05 |
| POYO | 0.90 ± 0.11 | 0.40 ± 1.27 | 0.95 ± 0.12 | **0.86 ± 0.18** |
| NEDS | 0.90 ± 0.09 | 0.62 ± 0.29 | 0.95 ± 0.03 | 0.82 ± 0.09 |
| **LAND** | **0.95 ± 0.04** | **0.73 ± 0.27** | **0.98 ± 0.00** | 0.72 ± 0.12 |

#### Cross-subject generalization

We transferred LAND in both directions using 80% of source windows and 20% of target-subject windows for adaptation, fine-tuning the target neural encoder, flow field, and prediction heads while retaining source-initialized behavioral dynamics. LAND achieved the highest held-out position and velocity *R*^2^ after transfer in both directions (Table 4). As visualized in Figure 5c, position decoding was near ceiling across methods, so differences in this metric should be interpreted cautiously. In contrast, transfer substantially improved velocity decoding from 0.85 to 0.94 for D→C and from 0.76 to 0.95 for C→D. The speed-ordered cyclic latent trajectories were retained after fine-tuning (Figure 5d), consistent with adaptation preserving learned cycling geometry while improving target-subject dynamics (see Supplementary for detailed data).

**Table 4:** Cross-subject held-out target decoding *R*^2^. Each entry reports *Target-only / Transfer*; higher is better.

| Method | D→C |  | C→D |  |
| --- | --- | --- | --- | --- |
|  | Position | Velocity | Position | Velocity |
| NDT2 | 0.92/0.91 | 0.87/0.87 | 0.92/0.88 | 0.90/0.89 |
| POYO | 0.87/0.90 | 0.82/0.86 | 0.89/0.92 | 0.86/0.88 |
| NEDS | 0.93/0.97 | 0.81/0.82 | 0.94/0.94 | 0.84/0.85 |
| CEBRA | 0.86/0.94 | 0.79/0.86 | 0.93/0.93 | 0.84/0.88 |
| **LAND** | **0.99/0.99** | **0.85/0.94** | **0.95/0.98** | **0.76/0.95** |

#### Ablation Studies

We quantified each LAND component across cross-speed decoding and limited-calibration transfer (Table 5). Entries are cross-domain *R*^2^, averaged over elbow sides for the epidural BCI, monkeys for NHP, and transfer directions. End-to-end training without auxiliary data sharply reduced cross-speed performance, especially epidural BCI velocity (0.114 versus 0.208). Removing auxiliary data caused the most consistent degradation, underscoring the importance of the auxiliary behavioral prior (details in the Supplementary).

**Table 5:** Ablation results (*R*^2^ for cross-domain decoding). EA/EV: Epidural BCI angle/velocity, averaged across sides; NP/NV: NHPs position/velocity, averaged across monkeys. Limb and Subj average both transfer directions.

| Setting | Speed |  |  |  | Limb |  | Subject |  |
| --- | --- | --- | --- | --- | --- | --- | --- | --- |
|  | EA | EV | NP | NV | EA | EV | NP | NV |
| E2E Training | .283 | .114 | .916 | .668 | — | — | — | — |
| w/o auxiliary data | .088 | .043 | .880 | .533 | .332 | .171 | .985 | .950 |
| w/o flow alignment | .330 | .142 | .925 | .648 | .372 | .214 | .985 | .940 |
| Embedding MSE align. | **.339** | .154 | .923 | .686 | .395 | .235 | .979 | .928 |
| w/o factor alignment | .205 | .081 | .917 | .667 | .378 | .209 | .986 | **.971** |
| w/o latent diversity | .315 | .160 | .927 | .683 | .395 | .226 | .984 | .917 |
| Unfrozen manifold | — | — | — | — | .313 | .192 | .983 | .943 |
| **LAND** | **.337** | **.208** | **.962** | **.723** | **.403** | **.252** | **.987** | .945 |

Direct embedding-MSE alignment was lower than LAND for velocity, NHP, and transfer metrics, suggesting a benefit from flow-based transport. The remaining ablations support complementary roles for the model components. Removing flow or latent-factor alignment reduced cross-speed performance in most settings, whereas omitting latent diversity degraded transfer in most settings. Unfreezing the behavioral manifold also reduced transfer performance. Although individual variants achieve the best performance on isolated metrics, full LAND provides the strongest overall cross-domain performance.

## Discussions and Conclusions

In this work, we introduce LAND, a framework for BCI movement decoding that leverages unpaired behavioral data. Rather than equating neural and behavioral embeddings, LAND learns a flow-based transport to a behavioral dynamical manifold and processes both paths with a shared causal decoder. This design incorporates behavioral temporal structure while preserving neural-only inference. Across synthetic, epidural, and NHP recordings, LAND improved zero-shot cross-speed decoding and, with limited paired calibration, cross-limb and cross-subject transfer. The speed-modulated latent geometry and ablations support contributions from the behavioral prior, flow alignment, and shared dynamical manifold.

The clinical evaluation uses one participant, and the NHP analysis two subjects with simple kinematics; broader multisession, multi-degree-of-freedom, and long-term closedloop evaluations are needed to establish robustness. Auxiliary trajectories come from controlled synthetic distributions or pose-estimated behavior, whose coverage and quality can limit the learned prior. Future work should learn richer behavioral primitives from naturalistic video (Duran et al. 2024; Singh et al. 2021), quantify how prior mismatch affects transport, and adapt the behavioral manifold when task structure genuinely changes. These directions may extend LAND toward scalable and reliable neural interfaces.

## References

Azabou, M.; Arora, V.; Ganesh, V.; Mao, X.; Nachimuthu, S.; Mendelson, M.; Richards, B.; Perich, M.; Lajoie, G.; and Dyer, E. 2023. A unified, scalable framework for neural population decoding. Advances in Neural Information Processing Systems, 36: 44937–44956.

Churchland, M. M.; Cunningham, J. P.; Kaufman, M. T.; Foster, J. D.; Nuyujukian, P.; Ryu, S. I.; and Shenoy, K. V. 2012. Neural population dynamics during reaching. Nature, 487(7405): 51–56.

Churchland, M. M.; and Shenoy, K. V. 2024. Preparatory activity and the expansive null-space. Nature Reviews Neuroscience, 25(4): 213–236.

Duran, E.; Kocabas, M.; Choutas, V.; Fan, Z.; and Black, M. J. 2024. Hmp: Hand motion priors for pose and shape estimation from video. In Proceedings of the IEEE/CVF Winter Conference on Applications of Computer Vision, 6353–6363.

Gallego, J. A.; Perich, M. G.; Miller, L. E.; and Solla, S. A.2017. Neural manifolds for the control of movement. Neuron, 94(5): 978–984.

Glaser, J. I.; Benjamin, A. S.; Chowdhury, R. H.; Perich,M. G.; Miller, L. E.; and Kording, K. P. 2020. Machine learning for neural decoding. eneuro, 7(4).

Jurkiewicz, M. T.; Gaetz, W. C.; Bostan, A. C.; and Cheyne,D. 2006. Post-movement beta rebound is generated in motor cortex: evidence from neuromagnetic recordings. Neuroimage, 32(3): 1281–1289.

Kao, J. C.; Stavisky, S. D.; Sussillo, D.; Nuyujukian, P.; and Shenoy, K. V. 2014. Information systems opportunities in brain–machine interface decoders. Proceedings of the IEEE, 102(5): 666–682.

Kapoor, J.; Schulz, A.; Vetter, J.; Pei, F.; Gao, R.; and Macke,J. H. 2024. Latent diffusion for neural spiking data. Advances in Neural Information Processing Systems, 37: 118119–118154.

Karpowicz, B. M.; Ali, Y. H.; Wimalasena, L. N.; Sedler,A. R.; Keshtkaran, M. R.; Bodkin, K.; Ma, X.; Rubin, D. B.; Williams, Z. M.; Cash, S. S.; et al. 2025. Stabilizing brain-computer interfaces through alignment of latent dynamics. Nature Communications, 16(1): 4662.

Keshtkaran, M. R.; Sedler, A. R.; Chowdhury, R. H.; Tandon, R.; Basrai, D.; Nguyen, S. L.; Sohn, H.; Jazayeri, M.; Miller,L. E.; and Pandarinath, C. 2022. A large-scale neural network training framework for generalized estimation of single-trial population dynamics. Nature Methods, 19(12): 1572–1577.

Komeiji, S.; Mitsuhashi, T.; Iimura, Y.; Suzuki, H.; Sugano, H.; Shinoda, K.; and Tanaka, T. 2024. Feasibility of decoding covert speech in ECoG with a Transformer trained on overt speech. Scientific Reports, 14(1): 11491.

Le, T.; and Shlizerman, E. 2022. Stndt: Modeling neural population activity with spatiotemporal transformers. Advances in Neural Information Processing Systems, 35: 17926–17939.

Li, T.; Yan, Y.; Dou, F.; Song, W.; and Zhang, X. 2026. Cross-subject generalization for EEG decoding: a survey of deep learning methods. Progress in Biomedical Engineering, 8(2): 022013.

Lipman, Y.; Chen, R. T. Q.; Ben-Hamu, H.; Nickel, M.; and Le, M. 2023. Flow Matching for Generative Modeling. arXiv:2210.02747.

Liu, D.; Shan, Y.; Wei, P.; Li, W.; Xu, H.; Liang, F.; Liu, T.; Zhao, G.; and Hong, B. 2024. Reclaiming hand functions after complete spinal cord injury with epidural brain-computer interface. medRxiv, 2024–09.

Liu, X.; Gong, C.; and Liu, Q. 2022. Flow Straight and Fast: Learning to Generate and Transfer Data with Rectified Flow. arXiv:2209.03003.

Lorach, H.; Galvez, A.; Spagnolo, V.; Martel, F.; Karakas, S.; Intering, N.; Vat, M.; Faivre, O.; Harte, C.; Komi, S.; et al. 2023. Walking naturally after spinal cord injury using a brain–spine interface. Nature, 618(7963): 126–133.

Manning, J. R.; Jacobs, J.; Fried, I.; and Kahana, M. J. 2009. Broadband shifts in local field potential power spectra are correlated with single-neuron spiking in humans. Journal of Neuroscience, 29(43): 13613–13620.

Mathis, M. W.; and Mathis, A. 2025. Joint modelling of brain and behaviour dynamics with artificial intelligence. Nature Reviews Neuroscience, 1–14.

Merrick, C. M.; Dixon, T. C.; Breska, A.; Lin, J.; Chang, E. F.; King-Stephens, D.; Laxer, K. D.; Weber, P. B.; Carmena, J.; Thomas Knight, R.; and Ivry, R. B. 2022. Left hemisphere dominance for bilateral kinematic encoding in the human brain. eLife, 11: e69977.

Miller, K. J.; Leuthardt, E. C.; Schalk, G.; Rao, R. P.; Anderson, N. R.; Moran, D. W.; Miller, J. W.; and Ojemann, J. G. 2007. Spectral changes in cortical surface potentials during motor movement. Journal of Neuroscience, 27(9): 2424–2432.

Pandarinath, C.; O’Shea, D. J.; Collins, J.; Jozefowicz, R.; Stavisky, S. D.; Kao, J. C.; Trautmann, E. M.; Kaufman,M. T.; Ryu, S. I.; Hochberg, L. R.; et al. 2018. Inferring single-trial neural population dynamics using sequential auto-encoders. Nature methods, 15(10): 805–815.

Perkins, S. M.; Amematsro, E. A.; Cunningham, J.; Wang, Q.; and Churchland, M. M. 2025. An emerging view of neural geometry in motor cortex supports high-performance decoding. Elife, 12: RP89421.

Pfurtscheller, G.; and Da Silva, F. L. 1999. Event-related EEG/MEG synchronization and desynchronization: basic principles. Clinical neurophysiology, 110(11): 1842–1857.

Rabbani, Q.; Fifer, M. S.; Crone, N. E.; and Moro-Velazquez,L. 2025. Impact of Temporal Precision on Speech Synthesis Accuracy From Electrocorticographic Brain Signals. In ICASSP 2025-2025 IEEE International Conference on Acoustics, Speech and Signal Processing (ICASSP), 1–5. IEEE.

Ryoo, A. H.-W.; Krishna, N. H.; Mao, X.; Azabou, M.; Dyer,E. L.; Perich, M. G.; and Lajoie, G. 2026. Generalizable, real-time neural decoding with hybrid state-space models. Advances in Neural Information Processing Systems, 38: 51764–51791.

Sani, O. G.; Abbaspourazad, H.; Wong, Y. T.; Pesaran, B.; and Shanechi, M. M. 2021. Modeling behaviorally relevantneural dynamics enabled by preferential subspace identification. Nature neuroscience, 24(1): 140–149.

Sani, O. G.; Pesaran, B.; and Shanechi, M. M. 2024. Dissociative and prioritized modeling of behaviorally relevant neural dynamics using recurrent neural networks. Nature neuroscience, 27(10): 2033–2045.

Saxena, S.; Russo, A. A.; Cunningham, J.; and Churchland,M. M. 2022. Motor cortex activity across movement speeds is predicted by network-level strategies for generating muscle activity. Elife, 11: e67620.

Schneider, S.; Lee, J. H.; and Mathis, M. W. 2023. Learnable latent embeddings for joint behavioural and neural analysis. Nature, 617(7960): 360–368.

Sedler, A. R.; Versteeg, C.; and Pandarinath, C. 2023. Expressive architectures enhance interpretability of dynamics-based neural population models. Neurons, behavior, data analysis, and theory, 2023: 10–51628.

Shenoy, K. V.; Sahani, M.; and Churchland, M. M. 2013. Cortical control of arm movements: a dynamical systems perspective. Annual review of neuroscience, 36(1): 337–359.

Shibasaki, H.; Barrett, G.; Halliday, E.; and Halliday, A. 1980. Components of the movement-related cortical potential and their scalp topography. Electroencephalography and clinical neurophysiology, 49(3-4): 213–226.

Singh, S. H.; Peterson, S. M.; Rao, R. P.; and Brunton, B. W. 2021. Mining naturalistic human behaviors in long-term video and neural recordings. Journal of Neuroscience Methods, 358: 109199.

Song, Y.; Liu, B.; Li, X.; Shi, N.; Wang, Y.; and Gao, X. 2023. Decoding natural images from eeg for object recognition. arXiv preprint arXiv:2308.13234.

Sun, H.; Qi, Y.; Wu, X.; Zhu, J.; Zhang, J.; and Wang, Y. 2024. Decoding Joint-Level Hand Movements With Intracortical Neural Signals in a Human Brain–Computer Interface. IEEE Transactions on Cognitive and Developmental Systems, 16(6): 2100–2109.

Sussillo, D.; Stavisky, S. D.; Kao, J. C.; Ryu, S. I.; and Shenoy, K. V. 2016. Making brain–machine interfaces robust to future neural variability. Nature communications, 7(1): 13749.

Suzuki, S.; Nagashima, S.; Hirata, M.; and Sugiura, K. 2025. Cortical-SSM: A Deep State Space Model for EEG and ECoG Motor Imagery Decoding. arXiv preprint arXiv:2510.15371.

Vyas, S.; Golub, M. D.; Sussillo, D.; and Shenoy, K. V. 2020. Computation through neural population dynamics. Annual review of neuroscience, 43(1): 249–275.

Wu, W.; Black, M.; Gao, Y.; Serruya, M.; Shaikhouni, A.; Donoghue, J.; and Bienenstock, E. 2002. Neural decoding of cursor motion using a Kalman filter. Advances in neural information processing systems, 15.

Xie, Z.; Schwartz, O.; and Prasad, A. 2018. Decoding of finger trajectory from ECoG using deep learning. Journal of neural engineering, 15(3): 036009.

Xu, M.; Chen, Y.; Wang, D.; Wang, Y.; Zhang, L.; and Wei,X. 2021. Multi-objective optimization approach for channel selection and cross-subject generalization in RSVP-based BCIs. Journal of Neural Engineering, 18(4): 046076.

Yao, R.; Du, Z.; Liang, F.; Li, W.; and Hong, B. 2025. Dual-layer hand gestures decoding with wireless epidural braincomputer interface in a tetraplegia. In 2025 47th Annual International Conference of the IEEE Engineering in Medicine and Biology Society (EMBC), 1–6. IEEE.

Ye, J.; Collinger, J.; Wehbe, L.; and Gaunt, R. 2023. Neural data transformer 2: multi-context pretraining for neural spiking activity. Advances in Neural Information Processing Systems, 36: 80352–80374.

Zhang, Y.; Wang, Y.; Azabou, M.; Andre, A.; Wang, Z.; Lyu, H.; Laboratory, T. I. B.; Dyer, E.; Paninski, L.; and Hurwitz,C. 2025. Neural encoding and decoding at scale. ArXiv, arXiv–2504.

Zhu, Y.; Song, C.; Ouyang, W.; Yu, S.; and Huang, T. 2025. Neural Representational Consistency Emerges from Probabilistic Neural-Behavioral Representation Alignment. arXiv preprint arXiv:2505.04331.

